# Application of a simple device for “attracting” and “trapping” glioma cells in situ

**DOI:** 10.1101/2021.12.21.473650

**Authors:** Haimin Song, Runwei Yang, Runbin Lai, Jinglin Guo, Kaishu Li, Bowen Ni, Yawei Liu

## Abstract

Glioblastoma multiforme (GBM) is the most malignant adult brain tumor. The current adjuvant therapies for GBM are disappointing, which are based on cytotoxicity strategy. Thus, other ways should be explored to improve the curative effect. According to the strong invasive ability of GBM cells, we assume a new treatment strategy for GBM by developing a new cell trap device (CTD) with some kind of “attractive” medium loaded in it to attract and capture the tumor cells. The in vitro experiment showed that Hepatocyte Growth Factor (HGF) presented stronger chemotaxis on C6 and U87 cell line than the Epidermal Growth Factor (EGF) and Fibroblast Growth Factor (FGF). A simple in vitro CTD loaded with HGF was made and in vivo experiments results showed that HGF successfully attracted tumor cells from tumor bed in situ into the CTD. This study proposes the new strategy for GBM treatment of “attract and trap” tumor cells is proved to be feasible.

## Introduction

Glioblastoma multiform (GBM) is the most common primary malignant tumor in adults, accounting for about 50% of intracranial gliomas [1]. At present, the comprehensive treatments for GBM include the maximum safe range of total resection, radiotherapy, chemotherapy with temozolomide (TMZ) [2]. However, the median survival is only 14.9 months [3], and the 5-year survival rate is about 5.5% [4]. Although immunotherapy has made breakthrough progress in clinical research, there is no substantial progress in the immunotherapy for GBM due to the existence of a blood-cerebrospinal fluid barrier in the central nervous system and fewer types and numbers of immune cells than other parts of the body [5].

Basically, cytotoxic treatment is the main control strategy for GBM, trying to “siege and destroy” tumor cells in various ways; however, their effects are not satisfactory. GBM cells show diffuse infiltrating growth, which causes it is impossible to remove tumor cells completely by surgery [6–8]. Chemoresistance and radiation resistance of tumor cells also bring clinical challenges for GBM treatment [9] Moreover, the tumor cell invasiveness under this cytotoxic and anti-angiogenic therapy would be significantly enhanced [9–11], leading to treatment failure. As a result, the tumor relapsed quickly. Therefore, GBM treatment requires new strategies.

The strategy of “attract-and-kill” could be a solution. Van der Sanden B. proposed the concept of ecological traps for the treatment of gliomas in 2013, designing ecological traps to attract tumor cells and kill them [12]. An ecological trap is a novel or higher quality environment, where there is a signal that misleading an animal to leave its traditional habitat. An ecological trap can drive a local population to extinction [12, 13]. However, Studies on ecological trap in cancer have largely focused on the tumor microenvironment, rather than for therapeutic purposes [14]. Seib et al. designed a cell “trap” to simulate a bone marrow environment to attract tumor cells. This trap mimics the characteristics of bone marrow based on breast cancer and prostate cancer cells trending metastatic to bone marrow [15]. Azarin et al made a three-dimensional microporous scaffold from poly lactide-co-glycolic acid (PLGA) scaffold to catch metastatic breast cancer cells in vivo [16]. To date, there is little in vivo experimental support for these hypotheses. Hence, we present a new treatment concept for GBM, that is “attract-and-kill” strategy (Figure.1). In this study, we design a model with a small chamber filled chemoattractant to verify the feasibility of tumor cells migrating from the in vivo tissue to an in vitro device implanted in experimental rats (Figure. 2A, 2B).

**Figure1.**
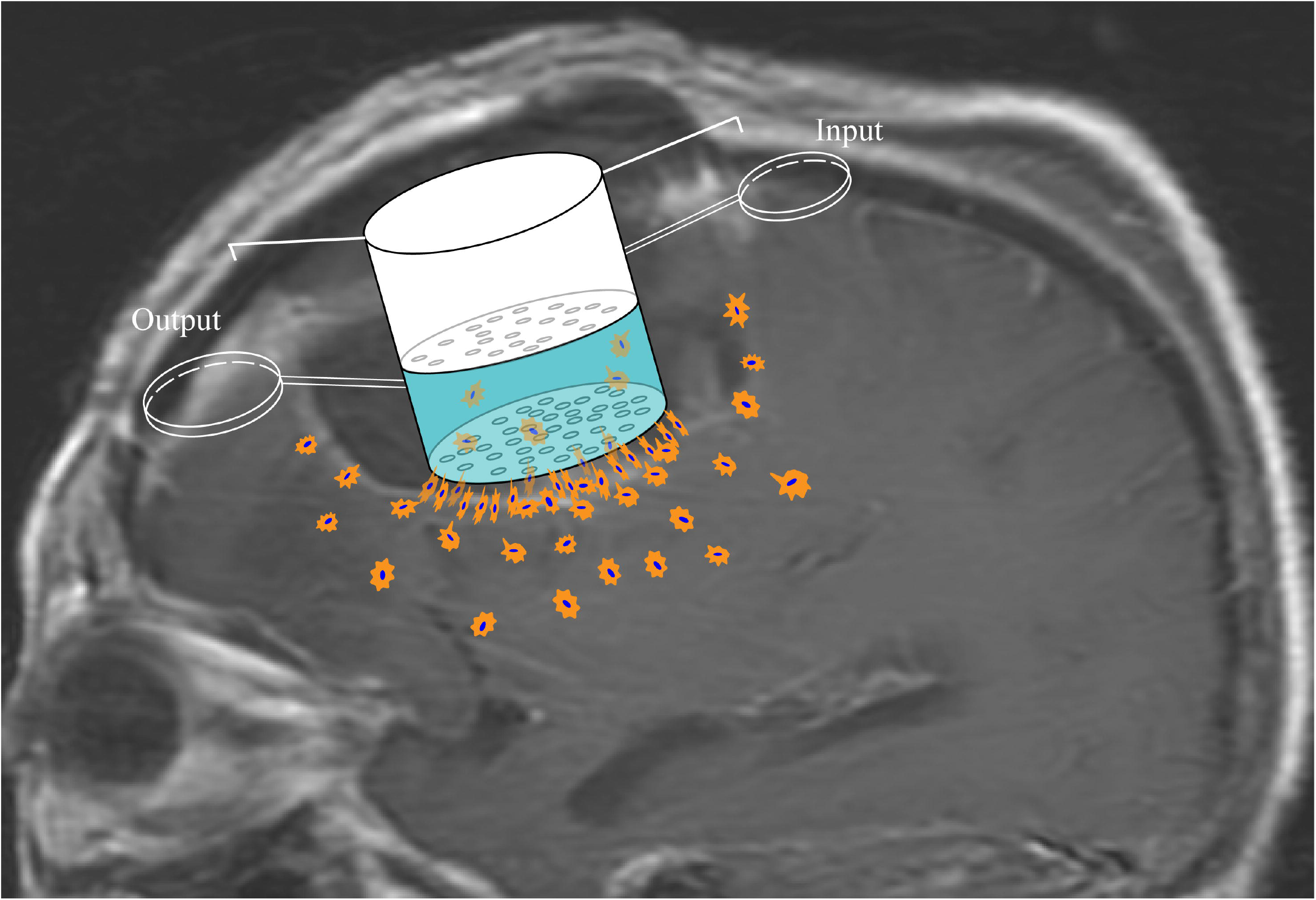
Treatment schematic of “attract-and-kill” for GBM. “Input” refers to attractive substance which contain cytokines, chemokines, chemoattractant and other substance induced cell migration. “output” means the site of egress of tumor cells from CTD.

**Figure2.**
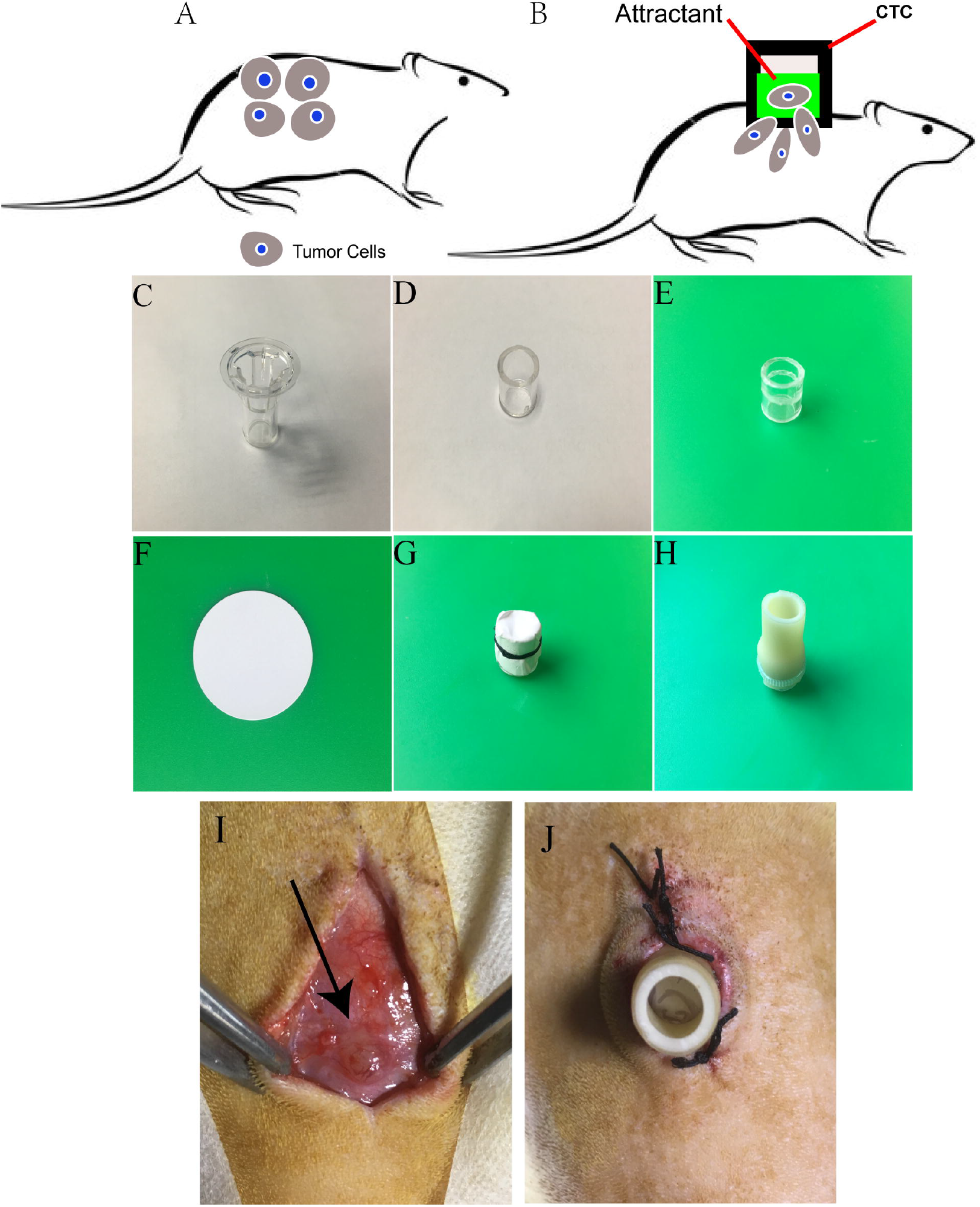
The animal experiment model of “attract-and-kill” subcutaneous glioma cells in rats. (A, B: CTD device carrying chemotactic substance was subsequently implanted into the subcutaneous, keeping the bottom membrane of the CTD clinging to the tumor bed and fixed. Tumor cells entered the device by the “attractive” chemokines; C: A transwell insert. D: The transwell was removed the upper half of opening; E: The insert was engraved 1 mm annular groove, F: A filter membrane; F, G: The opening of the instrument was covered with double filter membrane and fixed in the groove by the fourth thread, H: The soft tubes covered the opening and fasten the fixed catheter. I: incision of the subcutaneously exposed tumor, the black arrow indicated tumor; J: CTD basement membrane was fixed against the tumor and sutured to the skin. The part tube was exposed outside the skin.)

## Materials and methods

### Cell culture and lentiviral transduction

Rat glioma cell line C6 and human GBM cell line U87 (Cellular Library of the Culture Collection Committee of the Chinese Academy of Sciences) were cultured in Dulbecco’s Modified Eagle Medium (DMEM, Thermo Fisher Scientific, Gibco, #8118355) supplemented with 10% fetal bovine serum (FBS, Thermo Fisher Scientific, Gibco, #10099141) and 1% penicillin and streptomycin (Sigma-Aldrich, #P4333) in a 37°C incubator with 5% CO_2_. 5×10^5^ cells were plated into 6-well dishes in 2ml DMEM. The lentivirus (vigene biosciences, china) with 3×10^8^ TU in 10μl added to 6-well dishes per well when cells reached 70% confluence, and then changed with fresh medium 4 hours later. Stable C6 cells expressing green fluorescent protein (GFP)were selected with 10μM Puromycin (Solarbio, P8230, 5 μg/ml).

### Chemotaxis ability assays

The transwells with 8.0 μm prore polycarbonate membrane insert (Corning, #3422, USA) were used to detect the migratory ability of glioma cells. The polycarbonate membranes were coated with Matrigel™ (Corning, #356234, USA), 20 μg and 12.5 μg of matrigel were used for C6 and U87 cells respectively, and were incubated at 37°C for 1 hour before the glue solidified. The C6, U87 cells were cultured with DMEM medium containing 5% FBS for 6 hours at 80% fusion (the same below). Then digested into a monolayer cell suspension with 0.25% trypsin (containing EDTA), and resuspended by DMEM containing 0.2% bovine serum albumin (BSA). 100μl of 5×10^5^ cells suspension was placed on the upper layer of a cell culture insert with permeable membrane and a solution containing different cytokine by concentration gradient, HGF (R&D system, MN, #2207, USA), EGF (Peprotech, #AF-315-09, USA), or FGF (Peprotech, #100-18B, USA) is placed below the cell permeable membrane. C6 and U87 cells were incubated for 15 hours and 32 hours respectively. Then the chamber was fixed in 4% paraformaldehyde for 20 min, PBS washed twice, and stained with 0.2% crystal violet for 30 min. Wiped the cells not passing through the upper chamber with a cotton swab, dried them, and placed them under an inverted microscope (Lecia, German), randomly selected 5 fields to take photos and count the cells.

### Prepare the cell trapping chamber device (CTD)

the simple devices used in the animal experiment of this study were constructed by the commercialized Transwell inserts (Corning incorporated, ny14831, USA, 3422) with the bottom membrane aperture of 8.0 μm (Figure 2C): removed the upper ring, smoothed its opening (Fig. 2D), and carved an annular groove 3mm away from the opening without penetrating (Fig. 2E). Then, the circular filter membrane with a size of 25mm and a pore size of 0.15 μm (Fig. 2F) (Shanghai Xin ya Hydrophilic Hybrid Filter) was performed by ultraviolet radiation for 30 minutes in a biosafety cabinet. The collagen I (Solarbio, China, C8062) was prepared according to the three-dimensional cell culture formula and mixed with HGF (final concentration is 0.01nM, 0.1nM, 1nM and 10nM respectively). 200 μl collagen I with HGF was loaded into each device, the membrane surface at the bottom of the device was suspended for 20 minutes to coagulate the gel, then the double-layer filter membrane was used to cover the opening of the device, and the filter membrane was fixed at the groove with No. 4 surgical thread (Fig. 2G). Finally, sterile No. 24 tube was taken to cover the filter membrane and fixed with buckle (Fig. 2H). So far, the CTD was ready for use.

### SD (Sprague Dawley) rats

SD rats and rat derived glioma cells were used in this study. C6 glioma cell line was produced by Benda et al. [17] and it was reported to be the most similar to human glioma cells [18], so it is used for a variety of studies related to glioma biology, including tumor growth, invasion, migration, neovascularization [19–21]. The tumor formation rate of C6 in rat is 100% [22]. SD rats (purchased from the animal experimental center of Southern Medical University, production license number: SCXK (Yue) 2016-0041) of SPF grade adult male for 6-8 weeks, weighing 200-250g, were used in this study. The experimental rats were randomly divided into tumorigenic group and non-tumorigenic group. The tumorigenesis procedure was as follows: the rats were anaesthetized in the induction box of R450 type small animal anesthesia machine (Shenzhen Ruiwode Life Technology Co., Ltd., R580/C9M01-009), started with isoflurane (Shenzhen Rui Ward Life Technologies, Inc., 217180501) at a concentration of 4%, the air pump (R510-31S) flow rate was kept at 0.5 L/min and anesthesia-induced for 5 minutes. After the anesthesia was stable, the back of the rat was shaved to expose the skin between the front and rear limbs, and then disinfected with iodine (Shanghai Likang Disinfection Technology Co., Ltd., 20180713) and 3% hydrogen peroxide (Jiangxi Caoshanhu disinfection products Co., Ltd., 2018040). 5×10^6^ C6 cells were injected subcutaneously on both sides of the back and away from spine 0.5-1 cm. ALL animal experiments have been approved by the Nan Fang Hospital Ethics Committee.

### CTD implantation

The bulges of the dorsal mass were seen after 5 days of subcutaneous tumorigenesis. After anaesthetized, skin preparation and disinfection, an incision parallel to the spine was made to expose the tumor bed (Figure 2I). CTD was subsequently implanted, keeping the bottom membrane of the CTD clinging to the tumor bed and fixed. Then the skin was sutured with 4th silk. Part of the device was buried beneath the skin and part of which outside (Fig. 2J). After surgery, each rat was fed in a single cage.

### Multiphoton microscopy observation and processing

The CTDs were removed in 24 h and detected with multiphoton microscopy to find out whether were cells in there. The bottom film is sealed with blue butyl rubber (Blu-tack) (Bostik New Zealand Pty Ltd, 931049200459) water-sealed, and then CTD opening was covered with coverslip, also fixed and sealed with Blu-tack. Double distilled water was added to the dish and diffused over coverslip. The samples were scanned by multiphoton microscopy. Three fields of each sample were randomly selected and pictures were taken. The images were subjected to microscopic image analysis software Imaris (Imaris 9.0, BitPlane, BitPlane, Switzerland) for three-dimensional reconstruction and the background fluorescence intensity was adjusted to allow the cells to be fully exposed and photographed.

### Statistical analysis

Statistical analysis was conducted using SPSS 23.0 (IBMSPSS, Armonk, NY, USA). Data are presented as the mean±standard deviation of at least three independent experiments. Data performed homogeneity test of variance. If data meets homogeneity, Differences between multiple groups were analyzed using ANOVA, Otherwise, using nonparametric analysis. P < 0.05 was considered statistically significant. The two independent samples used a T-test for two groups comparison.

## Results

### HGF, EGF, FGF are chemotactic factors for C6, U87 in vitro, and HGF has the strongest chemotactic ability

The chemotactic ability of three cytokines HGF, EGF, FGF was tested through transwell experiments. We found that the chemotactic ability of HGF (0.1nM) was the strongest for C6 and U87 cells (Fig.3, S1, S2). U87 cells exhibited the strongest chemotactic response toward HGF at 0.1nM (p<0.001), EGF at 1nM (p<0.01), FGF at 50nM (p value is not statistically significant). C6 cells exhibited the strongest chemotactic response toward HGF at 0.1nM (p<0.001), EGF at 10nM (p<0.05), FGF at 10nM (p<0.01). (Figure 3, S1, S2). We examined the ability of three chemoattractant chemotaxis toward the two cell lines and found that HGF have the strongest chemotactic ability (p<0.05) (Figure.3). Therefore, we chose HGF as a chemotactic substance for animal experiments.

**Figure3.**
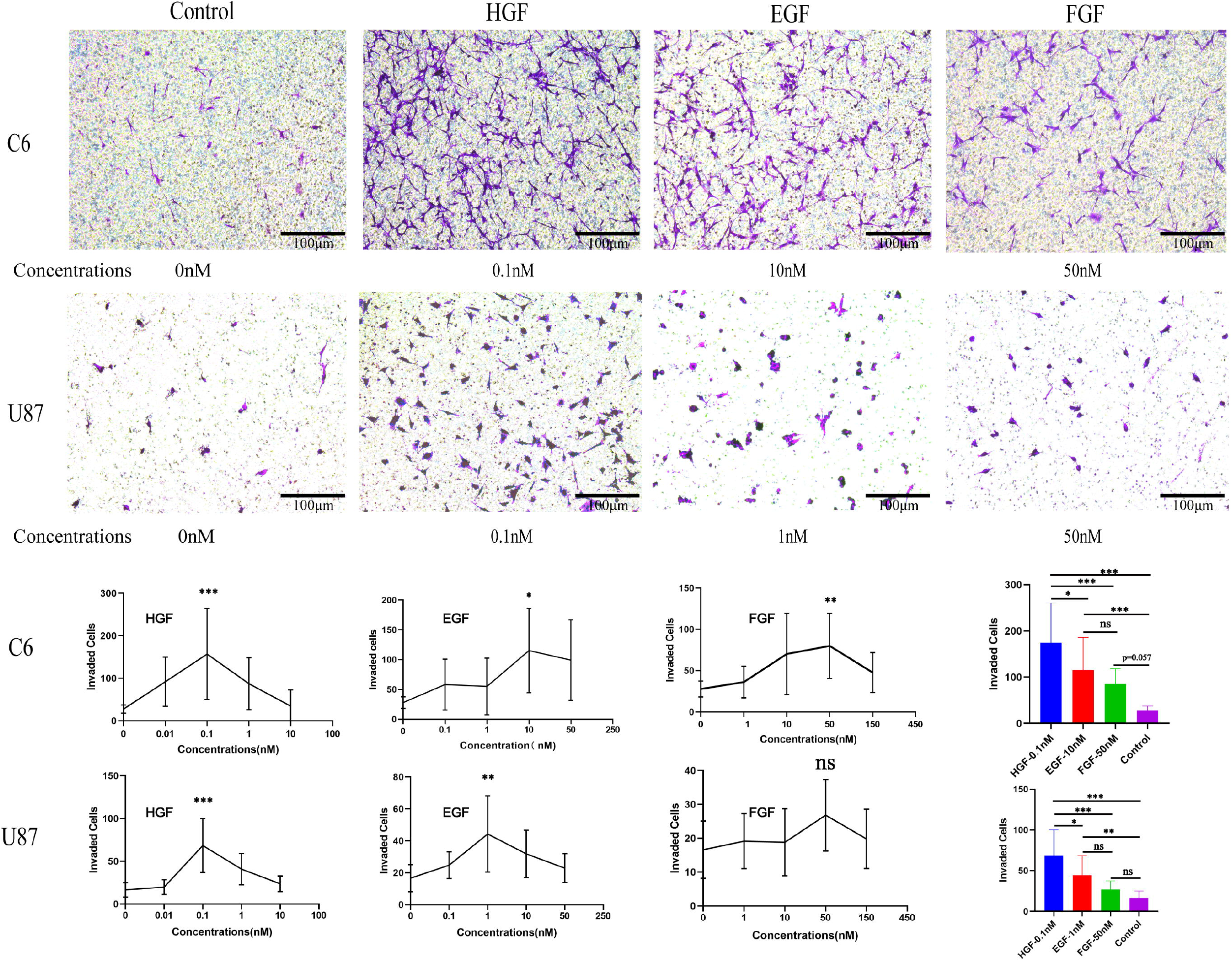
Chemotaxis of C6 and U87 cells toward HGF, EGF, FGF. * is P<0.05, ** is p<0.01, *** is p<0.001 and ns shows no statistically significant.

### HGF showed the ability of “attracting” tumor cells in vivo to extracorporeal CTD

To detect the presence of tumor cells in the CTD, GFP gene was transfected into C6 cells, so that the fluorescent intensities in the CTD could help to identify tumor cells. We proposed the following criteria to judge the fluorescent substance in CTD as C6 cells labeled by GFP. First strong fluorescent substances showed cell-like shape with size of about 10-20μm. Second, when the parameter of background fluorescence was weak or disappeared, the round or quasi-circular fluorescent substances still can be visible. Third, the fluorescent substance is present in the matrigel. Last, the matrigel presented invaded tracks around the fluorescent substance (Figure 4A, B, C).

**Figure 4.**
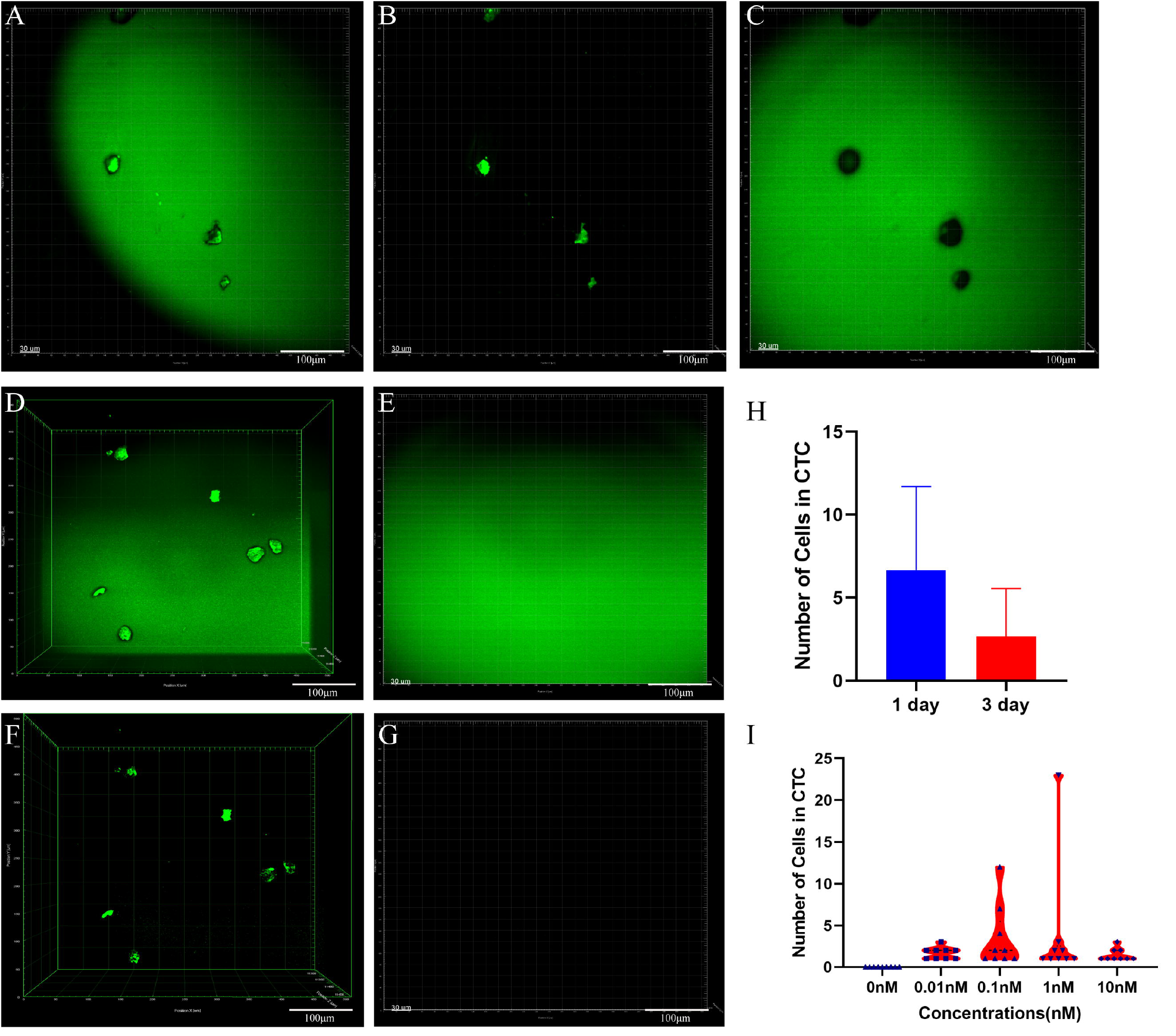
CTD MPLSM images (A, original images were taken by MPLSM; B, the cells; C, the matrix glue around the cells darkened by cell invasion. D, F showed cells in the CTD. E, G was the control group in which the tumorigenic group without HGF. H is the gradient time of chemotaxis in vivo; I was the violin plot of chemotactic effect of HGF at different concentrations).

We next demonstrated whether HGF has chemotaxis for C6 cells in vivo, as the CTD contains various concentrations of HGF matrigel (0.01nM, 0.1nM, 1nM, 10nM). Initially, we determined the optimal chemotaxis time of HGF in vivo, the result showed chemotactic effect 1 day was better than 3 days (Figure.4H). Therefore, we chose one day for animal experiment. After one day of contact between the membrane pore surface and the tumor, multiphoton microscope scanning found that HGF attracted tumor cells from the animal to the CTD, and had chemotactic ability at various concentrations. At a concentration of 0.1nM and 1nM, the trapped tumor cells were slightly more than that of other concentrations (Figure4. D, F, I). Despite the total number of trapped cells in various concentrations was few. Meanwhile, no fluorescent substance was found in the control group, which demonstrated that HGF had a chemotaxis effect for C6 cells in vivo (Figure 4E, G).

## Discussion

To validate the hypothesis that ecological trap can attract-and-kill glioma cells in animal models, a strong signal should be applied. Many factors can cause cell migration, such as neural precursor cells [23]and endogenous neural stem cells[24] under the action of the electric field. It was also noted that the magnetic field affected the migration of iron-labeled adipose stem cells [25]. But chemokines have been widely studied in Chemotaxis of tumor cells in vitro [26–28]. Ligand-derived factor (SDF-1) and matrix-derived factor receptor series (CXCR4, CXCR7) ligand receptors are applied to bone, blood vessels, and muscles researches because of chemotaxis [29–32]. One study in vivo was to penetrate the breast cancer tumor using the puncture needle, which contained EGF, EGF for the attraction of tumor cells and macrophages into the needle [33]. However, there are few investigations focus on GBM in vivo. Bacterial cellulose (BC) membrane loading with HSA was found to be a “trap” material. But HSA is very abundant in blood, which may limit its clinical application [34]. Therefor chemoattractant were screened for the experiment in this study. HGF is a better choice for in vivo experiments than EGF and FGF. HGF induce chemotactic migration of GBM cells to implanted CTD device and has no disturbance to local microenvironment. For a clinical application, the chemoattractant as signal should be easy to use and be refreshed. Thus, we designed a CTD filled with HGF mixed collagen I. The bottom of the CTD is a filter membrane with 8 μm pore diameter, which is pressed against the bed of tumor tissue when implanted into the animal. Tumor cells can crawl into the CTD through the membrane and be detected by multiphoton microscopy.

Another issue needs to be addressed is the detection of the tumor cells migrated into the CTD from in vivo residence. The collagen I gel in chamber is large, with very few or even no cells in it. Multiphoton microscopy based on a multiphoton absorbing dye allows for deep tissue imaging while maintaining a high resolution and thus offers a good choice for this study [35]. Glioma cells with green fluorescence that were transplanted into rats facilitate detection by multiphoton microscopy, provided they had entered the chambers.

The results of this study confirm that attract-and-kill is a feasible strategy for glioma therapy. The soft texture of intracranial tissue may have increased the possibility of its clinical application. However, several issues require improvement. Most importantly, the efficiency of the glioma cells migration into the cambers is very low. The first challenge is a more powerful chemotactic signal. HGF is acceptable, but is not sufficient. Next, the media carrying the chemotactic signal should be easy to be refreshed. We propose that the collagen I gel be replaced by some kinds of liquid media. Thus, the tumor cells are easy to be carried away and the chemotactic signal is easy to be refreshed.

## Conclusions

In this study, a simple device was designed and manufactured, which used “attractive” chemoattractant to “trap” tumor cells from the body to the extracorporeal CTD and helped to eliminate the tumor cells in vitro. This strategy provides a new direction for the comprehensive treatment of gliomas and other solid tumors.

## Availability of data and materials

The data used and/or analyzed in the present study are available from the corresponding author on reasonable request.

S1 (supplement fig1): Chemotaxis of C6 cells toward HGF, EGF, FGF at different concentrations.

S2 (supplement fig2) Chemotaxis of U87 cells toward HGF, EGF, FGF at different concentrations.

