## Supplementary figures and images for "Application of a simple device for “attracting” and “trapping” glioma cells in situ"

### figure-s1

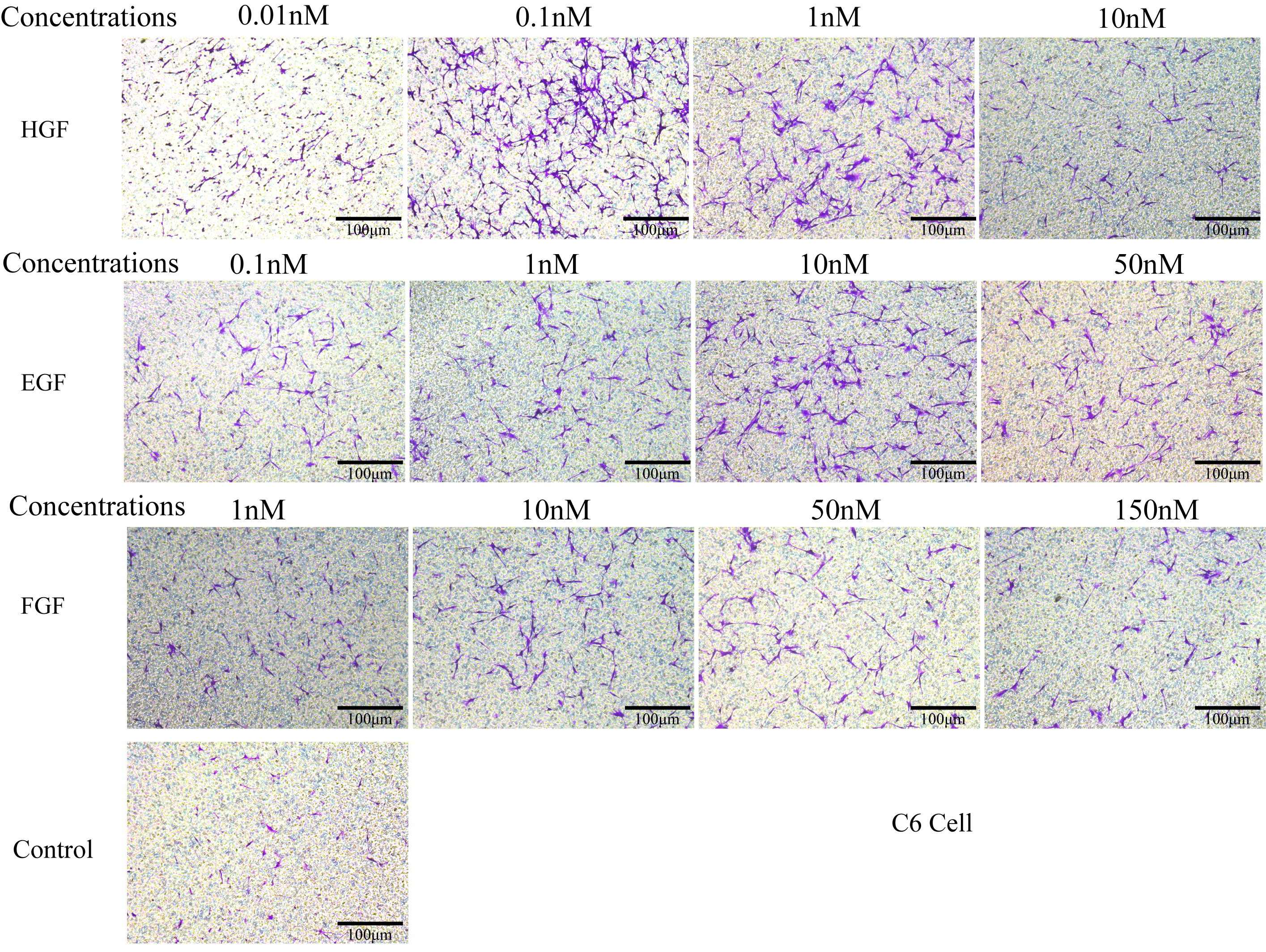

### figure-s2

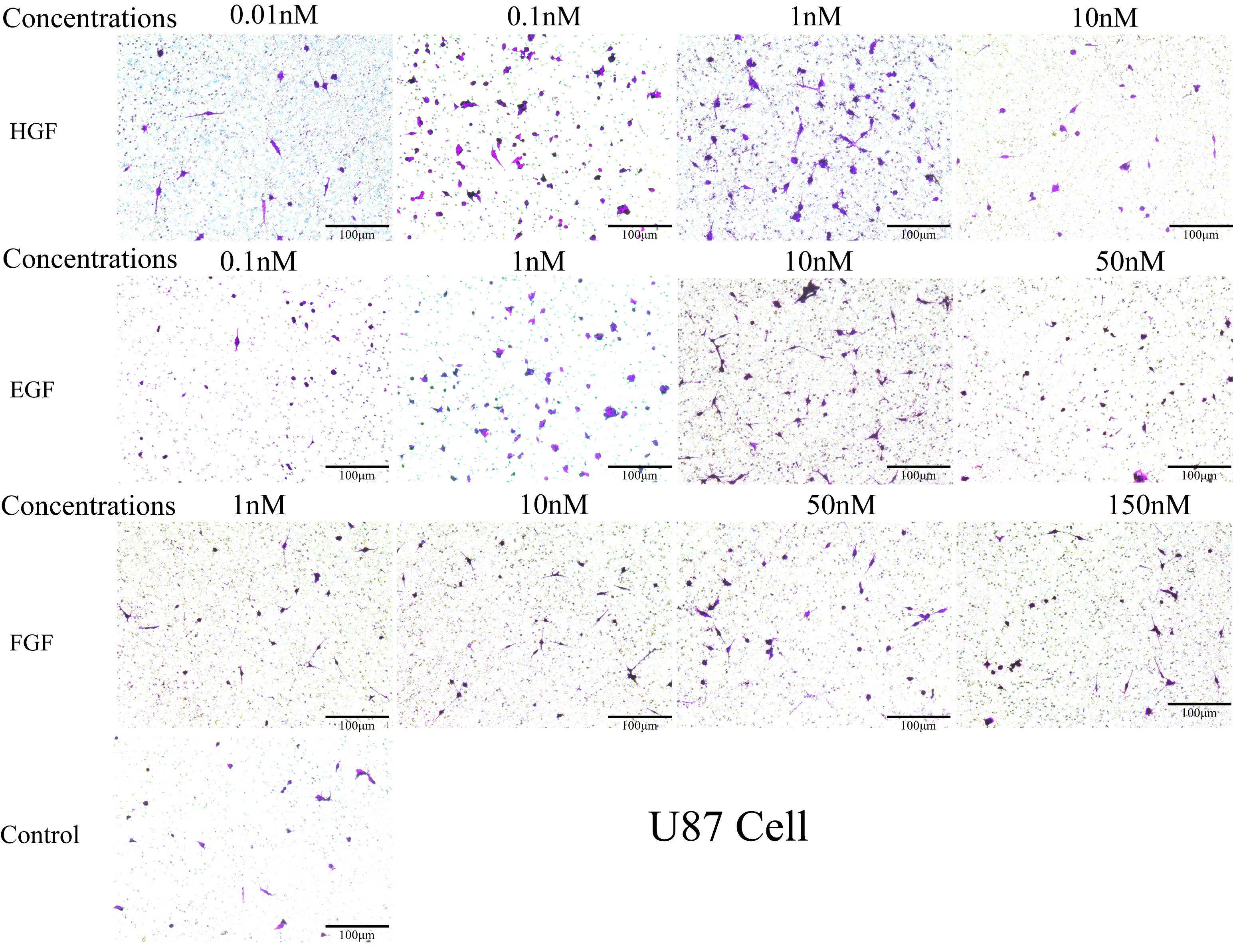
